# Root carbon interaction with soil minerals is dynamic, leaving a legacy of microbially-derived residues

**DOI:** 10.1101/2021.03.23.436628

**Authors:** Rachel A. Neurath, Jennifer Pett-Ridge, Ilexis Chu-Jacoby, Donald Herman, Thea Whitman, Peter Nico, Andrew S. Lipton, Jennifer Kyle, Malak M. Tfaily, Alison Thompson, Mary K. Firestone

## Abstract

1.

Minerals preserve the oldest most persistent soil carbon, and mineral characteristics appear to play a critical role in the formation of soil organic matter (SOM) associations. To test the hypothesis that carbon source and soil microorganisms also influence mineral-SOM associations, we incubated permeable minerals bags in soil microcosms with and without plants, in a ^13^CO_2_ labeling chamber. Mineral bags contained quartz, ferrihydrite, kaolinite, or native soil minerals isolated via density separation. Using ^13^C-NMR, FTICR-MS, and lipidomics, we traced plant-derived carbon onto minerals harvested from microcosms at three plant growth stages, characterizing total carbon, ^13^C enrichment, and SOM chemistry. While C accumulation was rapid and mineral-dependent, the accumulated amount was not significantly affected by the presence of plant roots. However, the rhizosphere did shape the chemistry of mineral-associated SOM. Minerals incubated in the rhizosphere were associated with a more diverse array of compounds with different C functional groups (carbonyl, aromatics, carbohydrates, lipids) than minerals incubated in a bulk soil control. These diverse rhizosphere-derived compounds may represent a “transient fraction” of mineral SOM, rapidly exchanging with mineral surfaces. Our results also suggest that many of the lipids which persist on minerals are microbially-derived with a large fraction of fungal lipids.

**Synopsis:** This study explores the interaction of rhizosphere carbon, minerals, and microbial influence on the fate of soil carbon.

**TOC:** 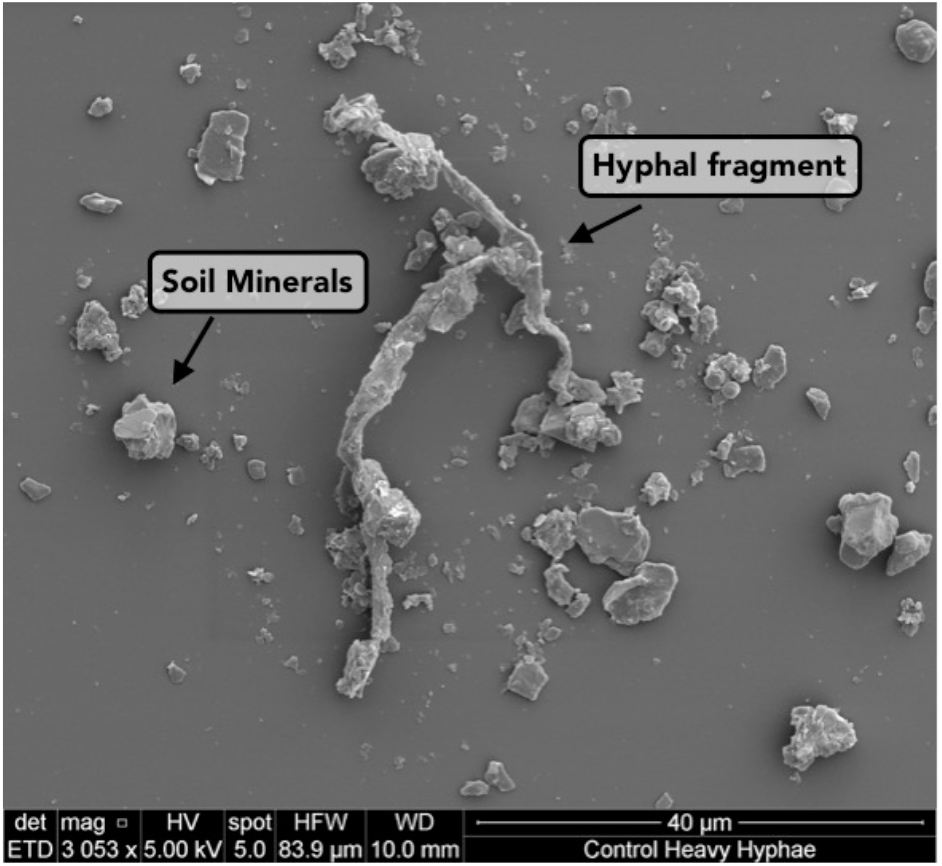

## 2. INTRODUCTION

Plant roots are the primary mediators of carbon (C) transfer into soil ^1-3^, allocating C captured from the atmosphere to the soil system ^4^. Root C is then transformed within the soil, where it may be respired back to the atmosphere or stored as soil organic matter (SOM) ^5^. Association with soil minerals offers a degree of protection for SOM ^6-11^, however the residence time of this SOM varies widely ^12^, due to a complex interplay of soil and microbial processes ^13-14^. Why some mineral-associated C persists and some does not is a central question in the field of soil C dynamics ^15^, with important implications for soil health, fertility, and microbial community ecology ^16-19^.

In grassland ecosystems, most surface soils are periodically or continually in the zone of root influence, or rhizosphere ^20^. As plant roots transfer organic compounds to the soil, the fate of this C is determined by (i) microbial community composition and activity ^19, 21- 23^, (ii) the chemical form of the C ^24^, (iii) where within the soil physical environment it is located ^25^, and (iv) soil mineral sorption capacity ^22, 26^. More than just a C source, growing roots physically and chemically alter the soil environment ^27-29^, with significant implications for the fate of soil C.

The rhizosphere is the nexus of plant-soil-microbe interactions ^30-31^. While plant growth provides a C source and stimulates microbial activity ^32-33^, root production of organic acids, such as oxalic acid, can release organo-metal complexes from mineral surfaces ^29^. Thus, roots influence both C association with minerals and also C loss. Microorganisms also interact with minerals, associating with mineral surfaces for protection from predation and desiccation, access to nutrients and energy sources, and as a platform for biofilm formation ^34-40^. Thus, it is now widely recognized that microbial necromass is a key part of mineral-associated SOM ^26, 41-43^. We hypothesize that the synergy of growing plant roots and rhizosphere microorganisms results in altered composition and quantity of mineral-associated SOM, an influence that takes place in the context of well-documented effects of mineral type on mineral-associated SOM quantity ^6^, composition ^44^, and persistence ^8, 24, 45-47^.

Here, we considered how biogeochemical differences between the rhizosphere and bulk soil might impact C persistence on minerals. As a zone of active C release and microbial activity, the rhizosphere is also characterized by both a lower degree of spatial heterogeneity and reduced microbial diversity^23^. Recent work by Lehmann *et al*. ^14^ suggests that functional complexity predicts soil carbon persistence. However, it is currently unknown whether the rhizosphere has higher or lower molecular functional diversity than bulk soil, but it is clearly more dynamic, with high diurnal ^48-49^ and seasonal ^33, 50^ temporal variability. Given this framework, we predict that rhizosphere SOM may turn over more rapidly than in the bulk soil.

In soil microcosms with living plants (*Avena barbata*), we incubated four mineral types in a ^13^CO_2_ growth chamber, allowing us to trace the fate of ^13^C-enriched plant-derived C. Our objectives were (1) to determine the influence of active roots on SOM association with mineral surfaces, (2) to determine the influence of mineral type on the quantity and composition of mineral-associated SOM, and (3) to explore the role of microorganisms in transformation of mineral-associated SOM. We found that while mineral type dictated the total quantity of mineral-associated SOM, the combined effect of mineral type and exposure to root growth defined the chemical composition of mineral-associated SOM. Rhizosphere SOM was compositionally more diverse than the bulk soil, but also appeared more transient, with higher turnover.

## 3. MATERIALS AND METHODS

### 3.1. Experimental Design

The study soil and plant type used in our soil microcosms were chosen to represent a typical Mediterranean-climate grassland ecosystem, with a fertile Alfisol soil and the naturalized slender wild oat grass *Avena barbata*. Both the soil and *A. barbata* seeds used in the soil microcosms were collected from Little Buck Field (38.992938° N, 123.067714° W), a managed pasture that is grazed by sheep, at the Hopland Research and Extension Center (HREC), Hopland, CA, in March, 2014, just before the start of the summer dry season. This field site is well characterized ^51^, and the microbial community has been studied extensively ^23, 32-33, 52-56^. The soil, a fine-loamy, mixed, active, mesic Typic Haploxeralf ^51^, was sieved to <2 mm in the field, dried to 1.1 volumetric water content (VWC), and stored at 4°C. Field soil had a field bulk density of 1.2 g-cm^-^3, a pH of 5.4 ± 0.03 (n = 10), and 23.3 ± 2.3 mg-g^-1^ (n = 3) total carbon (C). *A*. barbata seeds were collected in the field, dried, and scarified before germination.

Microcosms (2.9 × 11.5 × 25.3 cm) (n=30) were filled with soil to field bulk density (1.2 g-cm^-3^) (**SI Figure 1**), with 5 microcosms per treatment (bulk and rhizosphere) for each time point (1, 2, and 2.5 months). Before filling the microcosms, perforated tubing was inserted along one face of the microcosm (**SI Figure 1a**) to allow for even watering; in 12 microcosms, a volumetric water content (VWC) sensor (Decagon, EC5) was buried at approximately 10 cm depth and VWC was maintained at 14%, the average for Hopland soil during the growing season. Along the opposite face of the microcosm, a 2 mm width sidecar panel was inserted to keep that area free of soil and roots. We had two treatment types: (1) a *rhizosphere treatment*, with actively growing *A. barbata* planted in the soil microcosm, and (2) a *bulk soil treatment*, which was left unplanted. For the rhizosphere treatment, four germinated *A. barbata* seedlings were planted in each microcosm, simulating field density (**SI Figure 1b**). Microcosms were then tilted at 45°, with the sidecar facing downward, for one month (**SI Figure 1c-d**). Due to geotropism, roots grew along the sidecar panel face (**SI Figure 1e**).

After one month of plant growth, microcosms were opened along the sidecar face and the sidecar panel was removed, leaving a 2 mm gap. Nylon mesh bags (18 µm mesh, 5 × 5 × 0.2 cm dimensions) filled to field bulk density with one of four mineral types were placed in this gap, directly adjacent to the actively growing *A. barbata* roots, and then packed with soil. These minerals included: quartz sand (“Quartz”), ferrihydrite-coated quartz sand (“Ferrihydrite”), 50:50 by mass mixture kaolinite and quartz sand (“Kaolinite”), or the heavy density fraction (>1.75 g-cm^-1^) (“Native Minerals”) of Hopland soil (**Figure 1, SI Figure 1f**). Mineral preparation is described in the **SI Methods**. The specimen minerals added reflect the dominant mineral types present at our field site (determined by XRD, see **SI Methods**) and were selected to represent a range of surface area and reactivity, from quartz with low surface area and low reactivity to kaolinite with high surface area and moderate surface reactivity to ferrihydrite, an amorphous iron oxy-hydroxide, with moderate surface area and high surface reactivity (**SI Table 1**). The Native Minerals were density fractionated from field soil. We mixed kaolinite with quartz to prevent formation of large kaolinite aggregates and allow more permeability than pure kaolinite alone.

**Figure 1.**
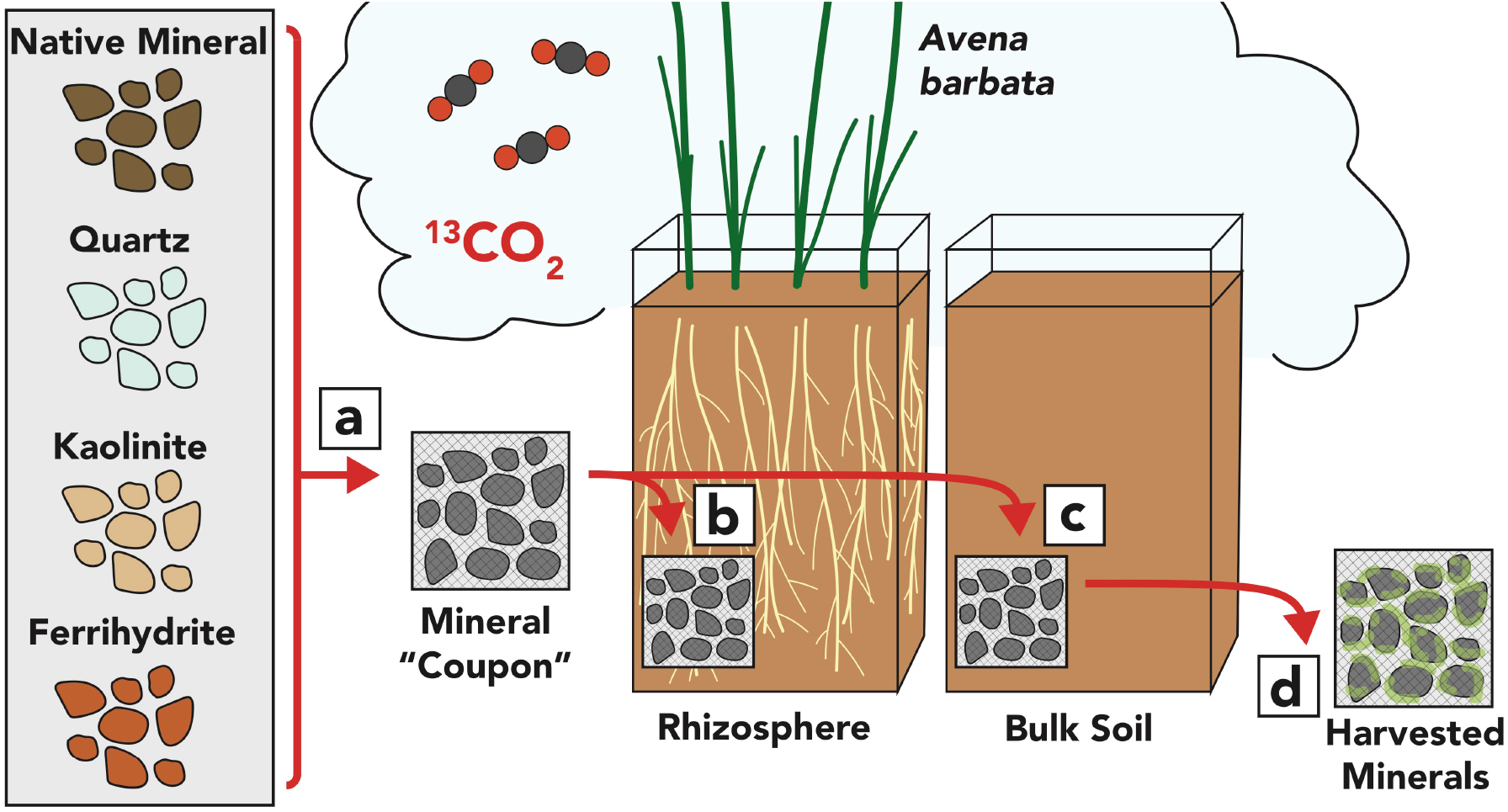
Experimental design for a rhizosphere mineral incubation in a Mediterranean-climate grassland soil from Hopland CA. Four mineral types, (1) “Native Minerals” (density fractionated from the soil), (2) “Quartz” (quartz sand), (3) “Kaolinite” (50:50 mix by volume of kaolinite and quartz sand), and (4) “Ferrihydrite” (ferrihydrite coated quartz sand), were placed in nylon mesh bags and placed in soil microcosms with growing *Avena barbata* plants (rhizosphere treatment) or no plants (bulk treatment). All microcosms were incubated in a 99 atom% ^13^CO_2_ labeling chamber. Microcosms were destructively harvested and mineral bags collected after 1, 2, and 2.5 months of incubation.

Mineral bags were harvested after 1, 2, and 2.5 months incubation, at which point *A. barbata* had reached senescence (**Figure 1, SI Figure 1g**). At harvest, both minerals and bulk soil were weighed, dried (65°C), and weighed again to obtain soil moisture content. The dried minerals and soil were stored at 4°C until analysis.

### 3.2. Total C and Isotope Ratio Mass Spectrometry

To measure total C accumulation on mineral surfaces, in bulk soil, and in root tissues, we used Elemental Analysis (EA) coupled to an IsoPrime 100 Isotope Ratio Mass Spectrometer (IRMS) (Isoprime Ltd, Cheadle Hulme, UK) to simultaneously measure total C and ^13^C enrichment in dried and ground minerals, bulk soil and roots. Peach leaf standards (NIST SRM 1547) were run to ensure accuracy and technical replicates for precision.

### 3.3. Scanning Electron Microscopy and BET Surface Area

Minerals were imaged with Scanning Transmission Electron Microscopy (SEM) at Lawrence Livermore National Laboratory (FEI, Inspect F) before and after incubation in the soil microcosms. Mineral samples were mounted and gold-coated and then images were acquired at 5 kV energy. The surface area of the minerals was determined before and after incubation by Brunauer-Emmett-Teller (BET) analysis with N_2_ gas at Lawrence Berkeley National Laboratory.

### 3.4. ^13^C-Nuclear Magnetic Resonance Mass Spectrometry

Major chemical classes of mineral-associated organic matter were identified by ^13^C-Nuclear Magnetic Resonance Mass Spectrometry. Finely ground samples were packed in a 5 mm optical density ceramic rotor and analyzed by 500 MHz solid-state ^13^C-NMR (Oxford) at the Environmental Molecular Sciences Laboratory (EMSL) at the Pacific Northwest National Laboratory (PNNL). Standards of potassium bromide and adamantine were run at 5,000 Hz. A soil standard, “Pahokee Peat” (IHSS, Standard Sample), collected from Florida and sieved to <53 μm, was also analyzed. The low C concentrations in our samples required long ^13^C-NMR experiments, thus we ran pooled samples (n=3) for each mineral type rather than replicates. Due to the interference of iron in the ferrihydrite sample and a low signal-to-noise ratio in the quartz sample, we were unable to obtain suitable ^13^C-NMR spectra from those mineral types (**SI Figure 4)**.

### 3.5. Fourier Transform Ion Cyclotron Resonance Mass Spectrometry

FTICR-MS and lipidomics (see section 3.6. below) analysis of mineral samples extracted with water (FTICR-MS) and then chloroform/methanol (lipidomics ^57^) was conducted at EMSL. Analyses were conducted on dried mineral samples from the 2-month time point. We ran 3 biological replicates of each sample.

FTICR-MS analysis is discussed in detail in SI Methods. Briefly, samples were dried and extracted with MeOH for chemical characterization on a 12T Bruker SolariX FTICR mass spectrometry, as previously described in Tfaily *et al*. (2017, 2018)^58-59^. Putative chemical formulas were assigned using Formularity software^60^. Compounds were plotted on van Krevelen diagrams based on their molar H:C ratios (*y*‐axis) and molar O:C ratios (*x*‐axis)^61^.

### 3.6. Lipidomics

Total lipid extracts were analyzed in both positive and negative modes, and lipids were fragmented using higher-energy collision dissociation and collision-induced dissociation. Confident lipid identifications were made using LIQUID (LIpid QUantitation and IDentification) ^62^. Aligned features were manually verified and peak apex intensity values were exported for statistical analysis. Identified lipids were selected by manually evaluating the MS/MS spectra for diagnostic and corresponding acyl chain fragment ions. In addition, the precursor isotopic profile, extracted ion chromatogram, and mass measurement error along with the elution time were evaluated. All LC-MS/MS runs were aligned and gap-filled using the identified lipid name, observed m/z, and the retention time using MZmine 2 ^63^. Each ionization type was aligned and gap-filled separately. Aligned features were manually verified and peak apex intensity values were exported for statistical analysis.

### 3.7. Sequential Extraction

To characterize the bonds between mineral surfaces and associated SOM, we performed a series of sequential extractions, adapted from Heckman *et al*. ^64^: (1) a water extraction was used to remove weakly mineral-associated soluble SOM (e.g. Van der Waals, dipole-dipole forces); (2) 0.1 M tetra-sodium pyrophosphate extraction ^65^ for organic-metal complexes and base soluble C; and (3) 0.25 M hydroxylamine ^65-66^ for non-crystalline and poorly crystalline iron and aluminum and associated C. We conducted sequential extractions on 3 biological replicates of each sample. For the sequential extractions, 40 mL extraction reagent was added to 3 g mineral sample, vortexed, shaken overnight, centrifuged and the supernatant decanted and passed through a 0.2 µm Whatman GX/D syringe filter. The residual minerals were rinsed with sterilized MilliQ water three times. The filtered supernatant was analyzed by ICP-MS (Perkin-Elmer Elan DRC II). The residual minerals were dried and analyzed for total C and ^13^C by EA and IRMS.

### 3.8. Mixing Model

To calculate the relative contribution of total C on mineral surfaces from the actively growing *A. barbata* plant, we used a mixing model (**Eq. 1**)^67^. In this model, the two pools of C were the *A. barbata* root and the bulk soil:

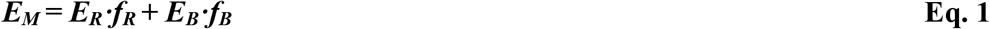

Where ***E***_***M***_ is the atom% enrichment of the mineral, ***E***_***R***_ is the atom% enrichment derived from the *A. barbata* root, ***f***_***R***_ is the fraction of C from the root, ***E***_***B***_ is the atom% enrichment derived from the bulk soil, and ***f***_***B***_ is the fraction of C from the bulk soil. In our equation, ***f***_***R***_ ***+ f***_***B***_ ***= 1***.

Bulk soil had a natural abundance ^13^C of 1.074 ± 0.001 atom%. Plant roots were pooled, ground, and analyzed by IRMS to calculate an average enrichment of 7.191 ± 0.398 SE atom% ^13^C.

### 3.9. Statistics

Carbon concentrations and ^13^C ratios are reported with standard error (n=5). We used a partially-nested mixed effects model according to Doncaster and Davey ^68^ to analyze the total C and IRMS results. We conducted analysis of variance (ANOVA) on these data, using the nlme package ^69^ in R v.3.5.1 (R Core Team, 2018) on log-transformed data to maintain assumptions of normality. To conduct pairwise comparisons, we used the emmeans package ^70^ in R. For FTICR-MS data, we performed non-metric multidimensional scaling (NMDS) analysis using presence/absence data and Jaccard distances to determine group differences, using the package ftmsRanalysis in R ^71^. To determine whether the differences illustrated in the NMDS plot were significant, we ran a permutational multivariate ANOVA (PERMANOVA) analysis in R using the adonis function in the vegan package^72^. We performed principal coordinate analysis (PCoA) with a Bray-Curtis dissimilarity matrix on the lipidomic data to determine sub-class differences in R using the vegan package ^72^. To determine whether the differences in the PCoA plot were significant, we ran a PERMANOVA analysis in R using the adonis function in the vegan package^72^.

## 4. RESULTS AND DISCUSSION

### 4.1. Mineral type controls total C accumulation on minerals

All specimen minerals (Quartz, Kaolinite, and Ferrihydrite) had no detectable C prior to incubation. Carbon accumulated on all specimen mineral types in both the rhizosphere and bulk soil treatments (**Figure 2 a, SI Figure 3**) and in SEM images, we observed visual evidence of C accumulation on the mineral surfaces (**SI Figure 2**). This C accumulation was rapid; the majority of C association with minerals occurred in the first month. While total C associated with kaolinite and quartz did not increase after 1 month of incubation, the enrichment of the C (atm%) associated with the minerals continued to increase until 2 to 2.5 months (Figure 2 a and b). Total C concentration varied significantly by mineral type (p<0.0001). When expressed as total C per gram mineral, the most C was associated with Ferrihydrite (1.3 ± 0.3 mg C-g^-1^ for rhizosphere at 2.5 months) (**Figure 2a**). However, when normalized to BET-measured surface area (**SI Table 1**), Quartz had the highest concentration of total C per square meter mineral surface (11.4 ± 1.6 mg C-m^-2^ for rhizosphere at 2.5 months) (**SI Figures 3-4**). This near reversal in trend reveals an important distinction in how total C measurements are presented. Here, we focus on total C per gram mineral because this better reflects how soil C stocks are expressed in most ecosystem assessments and model representation. We did not find a significant plant vs. bulk treatment effect for total C across all minerals (p=0.250) (**SI Figure 4**).

**Figure 2.**
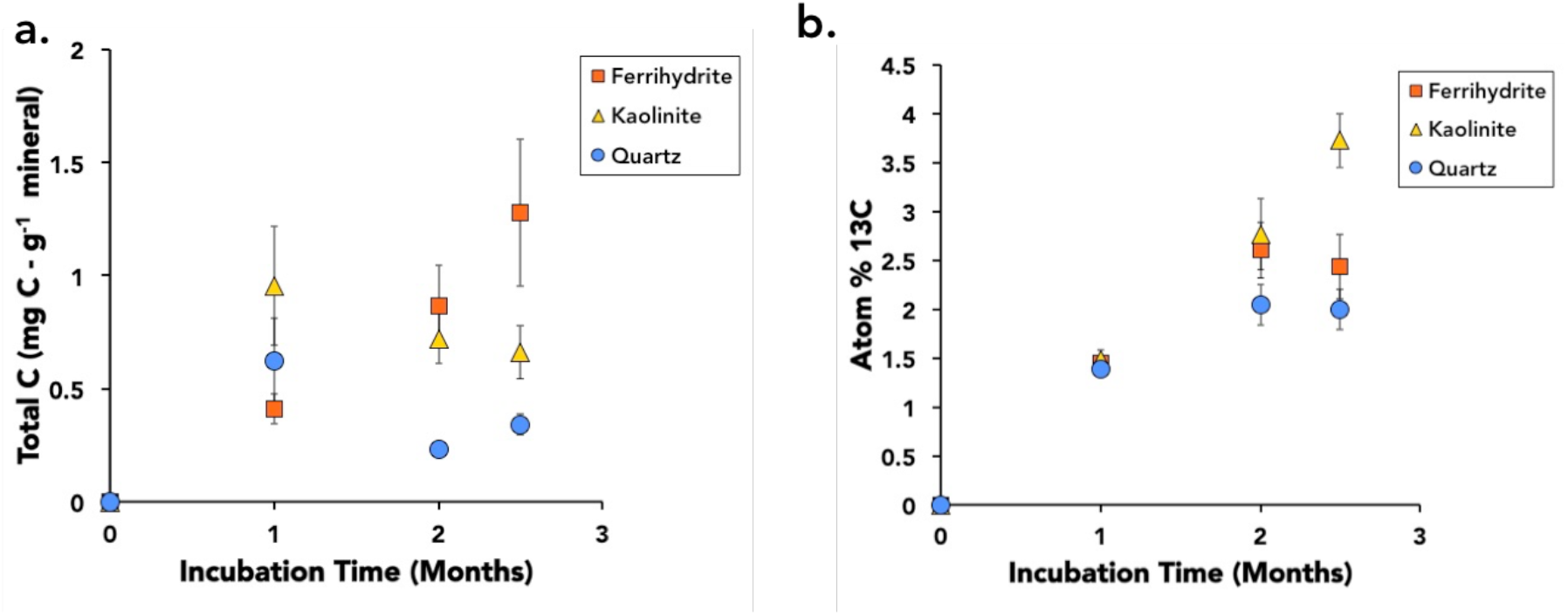
Total carbon (C) (a) and atom% ^13^C (b) of pure minerals: Ferrihydrite, Kaolinite, and Quartz, incubated in the rhizosphere of *Avena barbata* plants growing in a ^13^CO_2_ atmosphere. Panel (a) shows mineral total C accumulation over time. Panel (b) shows atom% ^13^C, reflecting mineral-associated C derived from the *A. barbata* roots. Error bars are standard error (N=5).

Carbon concentrations on specimen minerals were more than an order of magnitude lower than on the Native Minerals, with an average total C of 0.6 ± 0.1 mg C-g^-1^ for the specimen minerals in the rhizosphere versus 17.0 ± 1.0 mg C-g^-1^ for the Native Minerals (**SI Figure 4**). For our specimen minerals (Quartz, Ferrihydrite, and Kaolinite), we assume that at the end of the growing season we are still far from “carbon saturation” ^73-75^. Considering that the Native Minerals have likely resided in the soil for thousands of years or more ^76^, the relatively low specimen mineral C concentrations suggest that while C may accumulate rapidly on fresh mineral surfaces, the rate slows down over time.

Indeed, in the short 2.5 months of our study, we observed a linear rate of C accumulation on Ferrihydrite, whereas Quartz and Kaolinite minerals had rapid initial C accumulation followed by a leveling off (**Figure 2a**). The Native Minerals did not accumulate a significant amount of additional total C in either rhizosphere or bulk soil compared with their initial total C concentration (p = 0.101). Our study observes a single plant growing season, and therefore offers a critical snapshot of C accumulation and exchange on a mineral surface.

Rhizosphere-derived C contributed a significant portion of the C on mineral surfaces (**Table 1**) based on a ^13^C mixing model (**Eq. 1**). The percent rhizosphere-derived C significantly increased for all specimen minerals from 1 to 2 months (p < 0.001), and continued to increase significantly from 2 to 2.5 months for Kaolinite (**Figure 2b**). However, the rhizosphere treatment had no significant effect on the total amount of mineral-associated C. Thus, over the timescale of a single plant growing season, while there was evidence of plant-derived C on the minerals based on ^13^C tracing, the addition of root C to the microcosms did not result in a net increase in total mineral C, as compared to minerals that were incubated in bulk soil.

**Table 1.** **Percent C from *A. barbata* Rhizosphere**

| Time | Mineral | % Rhizosphere C |
| --- | --- | --- |
| **1 month** | Ferrihydrite | $4.89 \pm 5.5$ |
| | Kaolinite | $5.40 \pm 4.6$ |
| | Quartz | $1.44 \pm 3.4$ |
| **2 month** | Ferrihydrite | $24.0 \pm 5.5$ |
| | Kaolinite | $26.7 \pm 4.6$ |
| | Quartz | $14.7 \pm 3.4$ |
| | Native Mineral | $6.24 \pm 0.40$ |
| **2.5 month** | Ferrihydrite | $21.2 \pm 5.5$ |
| | Kaolinite | $42.6 \pm 4.6$ |
| | Quartz | $13.9 \pm 3.4$ |

A possible explanation for the absence of a rhizosphere effect on total C is that, while the total C stock remained the same, the fluxes of C were different. To illustrate, the stock of mineral associated C at time *t*, (***M***_***t***_), is equal to the stock of mineral associated C at time zero, (***M***_***0***_), plus the amount sorbed during time interval Δ*t*, (Δ***t·S***_***t***_), minus the amount desorbed during time interval Δ*t*, (Δ***t·D***_***t***_):

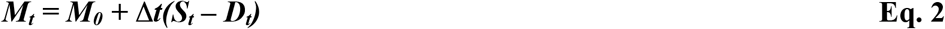

*where:*

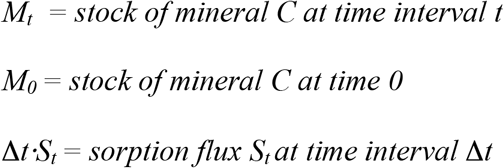

*where:*

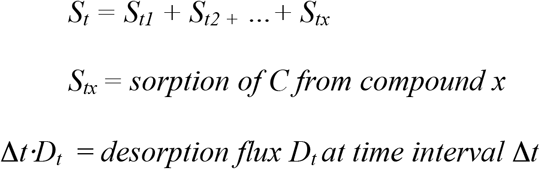

*where:*

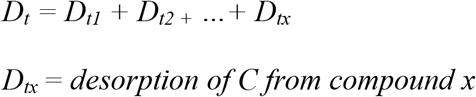

In our proposed equation, a modification of the classic mass-balance expression for net adsorption (See Eq. 8.8 ^77^), Δ*t·****S***_***t***_, is the aggregate sorption for each C compound that associates with the mineral, ***S***_***tx***_. Then Δ*t·****D***_***t***_, is the aggregate desorption for each C compound that associates with the mineral, ***D***_***tx***_. The ***S***_***tx***_ of a particular compound -for example, lipids -may not be the same as the ***D***_***tx***_ for that same compound; indeed, we believe they often are different. Thus, it is not only the rate at which compounds are sorbed and desorbed from the mineral, but also the composition of those compounds that matters, as we discuss further in section 3.4.

We hypothesize that in the rhizosphere, there is a larger supply of C^20, 23^, but there is also a faster turnover rate of mineral-associated C, which is consistent with Lehmann *et al*.’s ^14^ functional diversity framework, and the substantial literature on rhizosphere priming^78^ as well as recent work showing that more living microbes -as we would expect in the rhizosphere -can lead to less C retention^79^. For example, Kaolinite did not accumulate more total C after 1 month, and Quartz had a slight decline in total C (**Figure 2a**), while the atom% ^13^C of that C continued to increase (**Figure 2b**). We observed the same pattern when we calculated the percent contribution of ^13^C derived from the *Avena barbata* rhizosphere with our mixing model (**Eq. 1**). These results imply that there is active exchange of non-root derived C with new root derived C while maintaining the same total C concentration associated with the mineral. The chemistry of SOM associated with minerals incubated in the rhizosphere provides further evidence in support of our hypothesis, as discussed in section 4.2.

However, we cannot discount the possibility of a simpler explanation. If the rate of sorption of C to the mineral surface is not limited by the availability of C in the bulk soil, than it is possible that the rate of C exchange is the same as that in the rhizosphere. Further work is needed to test our hypothesis that mineral-associated C turnover is, indeed, faster in the rhizosphere than in bulk soil.

### 4.2. Mineral-associated SOM chemistry depends on mineral type and treatment

We used ^13^C-NMR to assess the molecular composition of mineral associated C. In contrast to the mineral total C accumulation, in the ^13^C-NMR spectra we observed a marked difference between rhizosphere and bulk soil treatments (**Figure 3**). While the ^13^C-NMR spectra for minerals in the rhizosphere show peaks across major organic carbon functional group (carbonyls, aromatics, carbohydrates, and lipids), the bulk soil treatment is defined by a single prominent lipid/aliphatic peak. Thus, while the rhizosphere treatment is characterized by a diverse array of carbon functional groups, the bulk soil treatment is distinct in its homogeneity. This suggests that the rhizosphere contributed a higher molecular diversity of organic C compounds than bulk soil.

**Figure 3.**
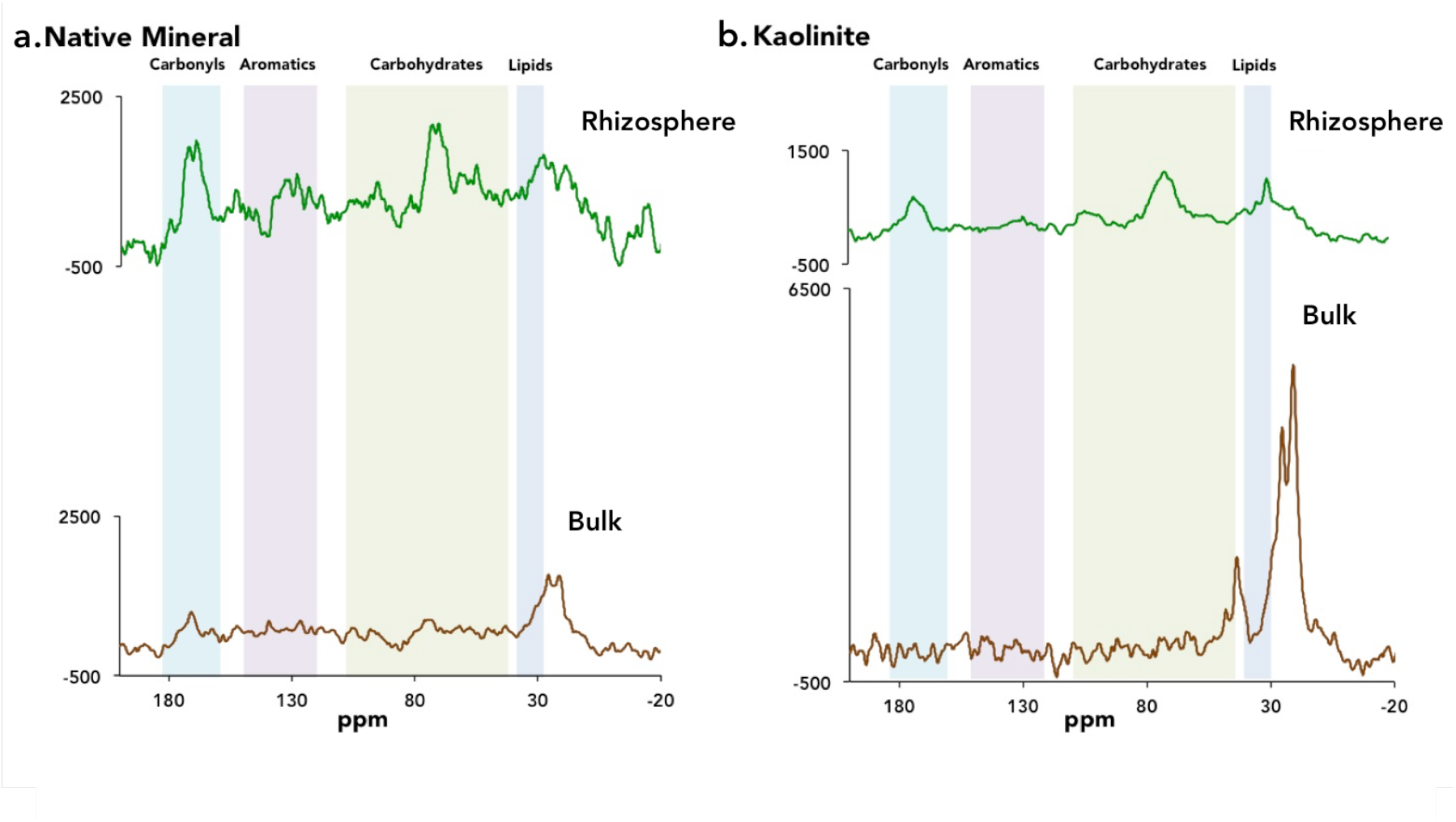
Solid-state ^13^C-NMR spectra for Native Minerals (separated via density fractionation) (a) and Kaolinite minerals (b), incubated in soil microcosms with rhizosphere and bulk soil treatments. Each spectra represents analysis of 3 pooled biological replicates. Peaks are clustered into broad organic carbon functional group categories: carbonyls, aromatics, carbohydrates, and lipids.

The striking contrast in organic C chemistry associated with the Native Minerals in the rhizosphere treatment versus those in the bulk soil suggests that the rhizosphere habitat dramatically altered the chemistry of mineral-associated SOM, even though only 6% of the C on the Native Minerals was directly derived from plant roots (based on our mixing model; **Table 1**). In the rhizosphere, roots are known to exude a wide array of small molecular weight compounds ^50^, promoting the growth of a phylogenetically distinct ^80^, and potentially chemically distinct microbial community. The ^13^C-NMR chemical signature of the rhizosphere Native Minerals suggests mineral-associated C is rapidly exchanging with a complex C pool comprised of rhizodeposits and associated organisms. By contrast, the carbon sources available in the bulk soil -which might include C from microbial and soil fauna products (*e*.*g*. extracellular polymeric substances (EPS) and microbial necromass) and dissolved organic C (DOC) – may be preferentially retained on the mineral surface such that only lipid/aliphatic residues remain. Since the chemistry of mineral-associated SOM is thought to be an important predictor of C persistence ^6-7, 45-46^, our results suggest that the relative proportion of minerals under the influence of actively growing roots versus those in a bulk soil setting may determine the overall molecular complexity of mineral-bound SOM. In addition, our finding that the Native Minerals acquired a unique rhizosphere chemical signature in a single plant growing season provides another line of support for our hypothesis that in the rhizosphere, mineral-associated C turns over faster.

We used FTICR-MS to measure the molecular composition of soluble SOM. These water-extractable compounds can be considered a “transient fraction” of mineral associated SOM: loosely associated compounds that are likely easily exchanged with soil pore water. Samples analyzed by FTICR-MS were significantly different by mineral type (p<0.001), but not treatment (rhizosphere versus bulk) (p>0.115) (**SI Figure 8**). The compounds detected by FTICR-MS (**SI Figure 9**) differ compositionally from those detected in the ^13^C-NMR (which represents the entire mineral-associated pool), suggesting that the transient, water-extractable fraction may represented only a subset of the total range of compounds present on the minerals. The extractability of these compounds appeared to depend on mineral type.

Given the apparent importance of mineral associated lipids observed in our ^13^C-NMR results, as well as in the literature^81^, we determined the fingerprint of lipid identity and distribution on our samples. We identified a total of 113 unique lipids. The distribution of unique classes of lipids was significantly different by mineral type (p<0.001) (**SI Figure 7**). The interaction of mineral type and rhizosphere versus bulk soil treatment was also significant (p<0.025) with a moderate effect of rhizosphere versus bulk soil treatment (p>0.063).

While differences in ionization potential prevent us from comparing between lipid categories, within a class we compared between mineral types and treatments (**Figure 4**). The Native Minerals had the highest lipid intensities for the majority of lipids observed in both rhizosphere and bulk soil treatments. Overall, Kaolinite had the second highest lipid intensities, with some triacylglyceride (TG) intensities higher than those on the Native Minerals. Overall, intensities of lipid classes associated with the specimen minerals types (Ferrihydrite, Kaolinite, and Quartz) were higher in the bulk soil treatment than the rhizosphere. This was particularly true of Ferrihydrite.

**Figure 4.**
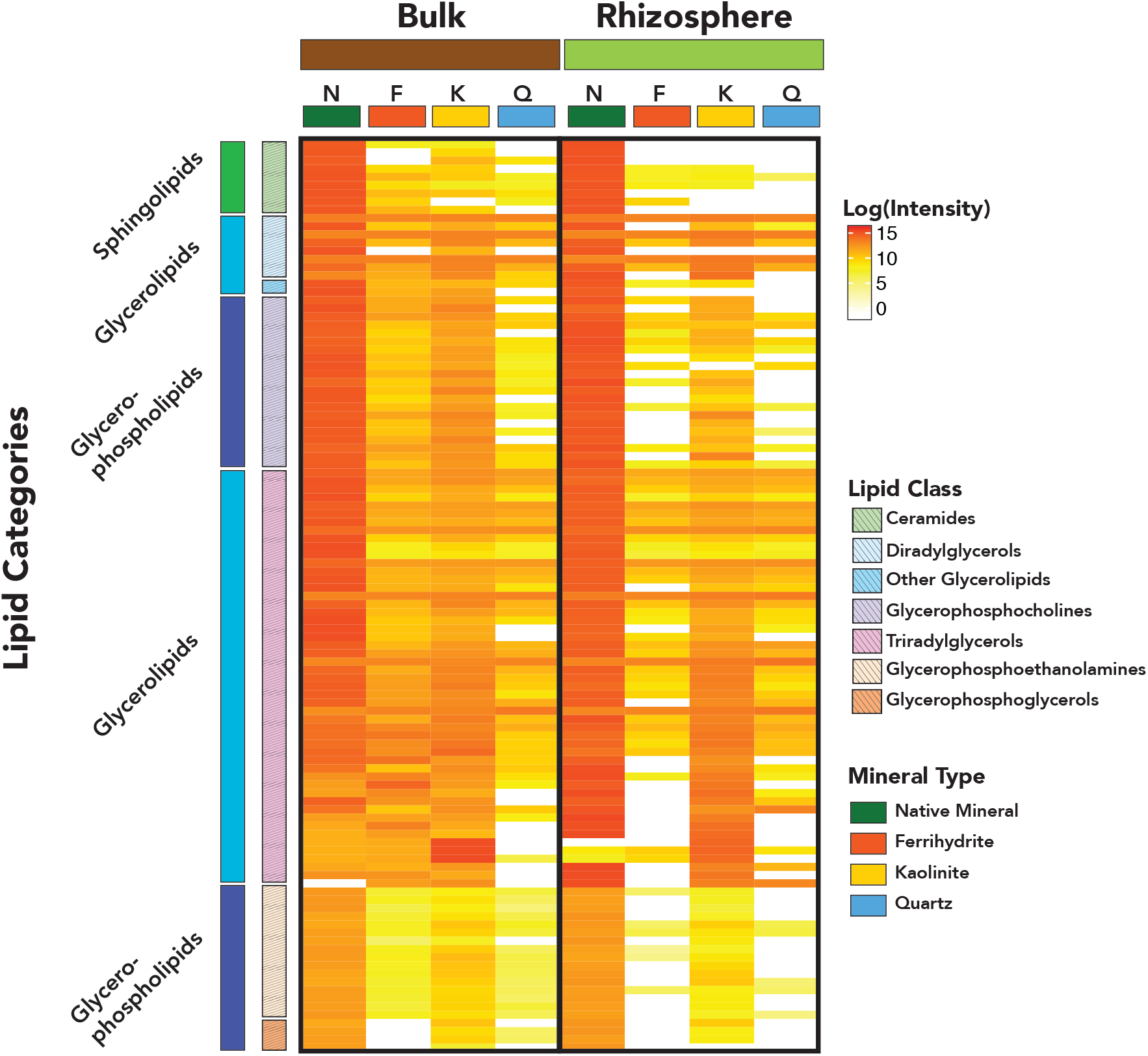
A heatmap comparison of log_10_ transformed lipid intensities for lipids identified in lipidomic analysis across treatment (bulk and rhizosphere) and mineral type (Native Mineral, Ferrihydrite, Kaolinite, and Quartz) (n=3). Due to differences in ionization potential, comparisons should only be made within lipid sub-class, rather than across broad lipid categories.

### 4.3. Microbial signatures in mineral-associated SOM chemistry

The largest class of lipids observed across mineral type and treatment was triacylglycerolipids (TG), with 59 total unique lipids. These storage lipids, which are highly abundant in fungi ^82^, are also found in plants and the bacterial genus *Actinomyces* ^83^. When we analyzed microbial colonization of the minerals from our study (published separately ^84^), *Actinobacteria* was the second most abundant phylum associated with the incubated minerals. The distribution of unique lipids within the TG class by mineral type was highly significantly different (p<0.001). Comparing lipid intensities, for many of the TG, Kaolinite had almost as high a lipid intensity as the Native Minerals, and for a few of the TGs, a higher intensity (**Figure 4**). Ferrihydrite also had high TG intensities in the bulk soil treatment. Thus, storage lipids may be an important component of mineral SOM.

We observed a diverse number of glycerophospholipids (GP), primarily of the classes diacylglycerophosphocholines (PC) and diacylglycerophophoethanolamines (PE). Highly abundant in bacteria^85^, these phospholipids are commonly structural components of cell membranes^86^. Many of the GP lipids we observed had odd numbers of C atoms (15, 17, or 19 C long fatty acids). Based on the structure of the Glycerophospholipids we identified, we believe that many were bacterial membrane lipids^87^.

Only 9 sphingolipids were observed, all of them in the ceramides and dihydroceramides sub-class. Ceramides are typically found in plants^82^, but are also present in most fungi and in a few anaerobic bacteria, such as *Bacteroides* spp^82, 85^, which are from one of the top ten bacterial phyla associated with the specimen minerals^84^. Interestingly, we observed more unique sphingolipids in the minerals from the bulk soil treatment than the rhizosphere treatment, which indicates that these lipids are more likely microbial than plant-derived (**Figure 4**).

Prior work on in our experimental system showed that mineral type shaped microbial association and assembly on mineral surfaces^84^. The distinct lipid signature we observed different mineral types may have been driven in part by the phylogenetically distinct microbial communities on these minerals. Chemical sorption may also play a role in determining which lipids end up on -or come off of -which mineral types. The microbial signature of mineral-associated lipids suggests that, if we know the soil mineral type, studying the microbial community composition and functional capacity may help in predictive modeling of mineral-associated SOM. Future research efforts examining the role of microbial ecophysiology on the fate of SOM may provide insights into the mechanisms through which soil microbes drive SOM persistence.

### 4.4. Towards predicting mineral-associated SOM persistence

Comparing the specimen minerals to Native Minerals allows for prediction of OM association over time. If we were to assume OM accumulates at a constant linear rate, in approximately 40 years, an average specimen mineral, initially free of detectable OM, would be coated in as much OM as the Native Minerals. However, we predict it would take far longer for a specimen mineral with no OM to accumulate the ∼ 16 mg C per g mineral we observed on Native Minerals because accumulation is not likely a linear, constant process^74-75^. In our results, we appear to be observing signals of a system in flux, as proposed in **Eq. 2**.

In the rhizosphere, an actively growing plant releases a diverse array of carbon substrates ^50, 88^. Our ^13^C-NMR results suggest that the rhizosphere leaves a distinct chemical fingerprint on mineral-associated OM. While some of these rhizosphere compounds may be sorbed directly from the root, we expect a large fraction of the mineral-associated OM is microbially derived. Indeed, the location of peaks in the rhizosphere mineral’s ^13^C-NMR spectra resemble those of *E. coli* cells, as analyzed by Wang *et al*. ^89^. Our prior work ^84^ reveals a diverse mineralosphere microbial community. Rhizosphere compounds could also be altered by abiotic processes prior to sorption. However, despite likely transformation of a portion of the rhizosphere-derived compounds, the rhizosphere treatment maintains a unique chemical signature that distinguishes it from the bulk soil treatment.

We see evidence that the unique chemical fingerprint of the rhizosphere may not persist, particularly in the comparison of the Native Minerals to Kaolinite. Prior to being field-collected, the Native Minerals were undoubtedly periodically rhizosphere influenced. However, the chemistry of Native Minerals incubated in the bulk soil closely resembles that of the Kaolinite mineral that was incubated for only two months in bulk soil. The similarity between the broad ^13^C-NMR chemistry of the Native Minerals and Kaolinite minerals in the bulk soil treatment suggest that, over time, many mineral-associated carbonyls and carbohydrates are either desorbed from the mineral surface or transformed, leaving behind mostly lipids.

The hydrophobic nature of lipids may lead to associations with mineral surface complexes ^90^ that are relatively difficult to desorb. In our system, lipids seemed to associate strongly with Kaolinite, which had the highest intensities of most lipid classes of the specimen minerals (**Figure 4**). Microbial assimilation of C compounds may result in transformation of those compounds to lipids as microbes synthesize membranes and storage lipids. We expect that over time, a larger portion of the mineral surface is occupied by microbial necromass, including microbial lipid residues.

While lipids may persist, sequential extractions indicate that a large portion of mineral-associated C was water extractable, and thus may readily exchange with the soil DOC pool. This fraction of extractable total C was strongly dependent on mineral type (**SI Figure 10**). Nearly all Quartz-associated C was removed via the water extraction (88% for rhizosphere, 100% for bulk soil), whereas a larger portion C remained on the Ferrihydrite and Kaolinite, highlighting the importance of stronger chemical associations between SOM and the mineral (*e*.*g*. ligand exchange) for C persistence (**SI Figure 10**)^77^. Interestingly, in the rhizosphere treatment, we saw that atom% ^13^C increased with each extraction for Kaolinite (**SI Figure 10**). This increase in atom% ^13^C of Kaolinite-associated C following an extraction indicates preferential persistence of *A. barbata-*derived C compounds. This suggests that chemistry of both the mineral surface and also the carbon substrate are important in controlling mineral-associated C persistence.

Our results provide evidence that SOM turnover is faster in rhizosphere than bulk soil. We also observe the potential importance of mineral type for predicting total SOM association with minerals. Prior work shows that short range order minerals, such as ferrihydrite, often have the oldest associated carbon ^6, 46-47, 91^. However, our study shows that mineral type influences on total C accumulation can be context-dependent (rhizosphere versus bulk soil). In the rhizosphere, Ferrihydrite had the highest total C, but in the bulk soil, Kaolinite had higher total C. Thus, mineral type alone may be insufficient to predict C accumulation and turnover. The presence or absence of root influence also appears to play a critical role in determining the chemistry and fate of mineral-associated C.

Our study emphasizes the importance of understanding plant-microbe-mineral interactions in concert to understand the mechanisms of C transformation and persistence in soil. We found critical differences between bulk soil and rhizosphere soils, which are likely particularly important in a seasonal system such as our model annual grassland ecosystem growing in a Mediterranean-type climate. While further work is needed to examine these systems over longer time scales, the results of this study provide a snapshot of C flow from plants to minerals over the course of a season as *A. barbata* grows and senesces. Carbon association with minerals is dynamic, particularly so in the rhizosphere. Based on our findings, we suggest that increasing C supply does not necessarily result in increased C association with minerals over a short timescale. Instead, differences in C source and environment impact mineral-associated C composition, which in turn could influence longer-term persistence.

## Acknowledgements

This research was supported by the US Department of Energy (DOE) Office of Science, Office of Biological and Environmental Research Genomic Science program under award DE-SC0016247 (to MKF) and awards SCW1589, SCW1421 and the LLNL Soil Microbiome SFA, SCW1632 (to JPR). RAN was supported by a Livermore Scholar Program fellowship at Lawrence Livermore National Laboratory and a National Science Foundation Doctoral Dissertation Improvement Grant, award 1601809. Part of this work was conducted at the Environmental Molecular Sciences Laboratory (grid.436923.9), a DOE Office of Science scientific user facility sponsored by the Department of Energy’s Office of Biological and Environmental Research and located at PNNL under contract DE-AC05-76RL01830. Work conducted at Lawrence Livermore National Laboratory was supported under the auspices of the U.S. DOE under Contract DE-AC52-07NA27344. Work conducted at Lawrence Berkeley National Laboratory was supported under Contract DE-AC02-05CH11231. Soil and plant collection were supported by the Hopland Research and Extension Center. Plants were grown and labeled at the University of California, Berkeley, Oxford Tract Greenhouse Facility. Sampling efforts and technical expertise were provided by Rina Estera-Molina.

Garrison Sposito provided invaluable insight and advice on the experimental design and interpretations.

