## Supplemental Information for "Root carbon interaction with soil minerals is dynamic, leaving a legacy of microbially-derived residues"

[a]Department of Environmental Science, Policy, and Management, University of California, Berkeley; [b]Biosciences Division, Lawrence Berkeley National Laboratory; [c]Physical and Life Sciences Directorate, Lawrence Livermore National Laboratory; [d]Department of Soil Science, University of Wisconsin, Madison; [e]Earth Sciences Division, Lawrence Berkeley National Laboratory; [f]Environmental Molecular Sciences Laboratory, Pacific Northwest National Laboratory; [g]Department of Environmental Science, University of Arizona

### SUPPORTING INFORMATION

#### 1. Supporting Methods

*1.1. X-Ray Diffraction*

To select minerals for the experiment that reflected the dominant mineral types found at Little Buck Field at HREC, we conducted X-Ray Diffraction (XRD) on the field soil (Panalytical X'Pert Pro diffractometer, Earth and Planetary Sciences Department, University of California, Berkeley). We analyzed finely ground bulk soil and then separated the clay fraction by suspension in deionized water and filtration. To resolve the clay mineralogy, we first analyzed the air-dried clay fraction. Next, we treated the clay fraction with ethylene glycol, air dried, and reanalyzed. Finally, we heated the clay fraction to 400°C, which allowed us to resolve the clay mineralogy. We also validated our lab-synthesized ferrihydrite-coated quartz using XRD. Peaks were analyzed using X'Pert Highscore Plus software and matched to reference peaks. We found that in the field soil, the dominant clay mineralogy was kaolinite. The dominant metal-oxide in the field soil was ferrihydrite, and we were also able to validate that our lab-synthesized ferrihydrite-coated quartz was ferrihydrite, and had not crystallized to goethite. The soil was also dominated by quartz. We also found feldspar, primarily albite, and mica, but these were not as dominant and so were not selected for the experiment.

*1.2. Mineral Preparation*

We prepared three pure mineral types for incubation in the soil microcosms, informed by the dominant mineralogy of Little Buck Field bulk soil (See **SI X-Ray Diffraction**). For simplicity, we refer to the four mineral types as: “Quartz” (quartz sand), “Kaolinite” (50:50 mix of kaolinite and quartz sand), “Ferrihydrite” (ferrihydrite-coated quartz sand), and “Native Minerals” (heavy fraction of density fractionated Hopland field soil). Quartz sand (Sigma-Aldrich, 50-70 mesh particle size, CAS 14808-60-7) was acid washed with 10% HCl, then rinsed with deionized water until reaching a neutral pH. Kaolinite was purchased from the Clay Minerals Society (KGa-2 Kaolin) and combined in a 50:50 by mass mixture with a subset of the acid-washed quartz sand. To synthesize the ferrihydrite-coated quartz, we used a modified method from Hansel *et al.* [92-93]. A solution of 34.4 mM $FeCl_3$, 6.87 mM $AlCl_3$, and 100 μm $Na_2SiO_3$ was mixed vigorously and the pH adjusted to 7.2-7.5 with the additional of 0.4 N NaOH, causing precipitation of the ferrihydrite. The supernatant was decanted and the ferrihydrite slurry was centrifuged at 3000 rpm. This process was repeated four times, with the centrifuge increased each time up to 8000 rpm. Centrifuge tubes with ferrihydrite slurry were sealed and left in a fume hood to age for one month. A subset of the acid-washed quartz sand was coated in the aged ferrihydrite slurry and mixed until homogenous, then dried. The ferrihydrite-coated quartz sand was washed with deionized water until it rinsed clear, then dried.

In addition to the three “pure” minerals prepared in the laboratory, we simulated “native” soil minerals by separating the heavy density fraction of soil collected at Little Buck Field (0-10cm). To separate the Native Minerals, we density fractionated air-dried soil samples. Following the density-fractionation method used in Pett-Ridge *et al.* [88], modified from Sollins *et al.* [94]we first separated the free light fraction (<1.75 g-cm$^3$). The remaining soil was sonicated to remove the occluded light fraction (<1.75 g-cm$^3$). We washed the remaining heavy fraction (>1.75 g-cm$^{-3}$) with deionized water three times and lyophilized.

*1.3. FTICR-MS Analysis*

Dissolved organic carbon in the water extract (soluble SOM) was concentrated on Bond Elut PPL cartridges (Agilent Technologies) that were pre-conditioned with HPLC-grade methanol as previously described by Dittmar *et al.* 2008[95]. After removing excess salts by flushing cartridges with 0.01 M HCl, cartridges were dried under pure $N_2$ gas and extracted with 2 mL of MeOH for SOC chemical characterization analysis on a 12T Bruker SolariX FTICR mass spectrometer by direct injection, as previously described in Tfaily et al. (2017, 2018)[58-59]. A standard Bruker ESI source was used to generate negatively charged molecular ions and then samples were introduced directly to the ESI source. The instrument was externally calibrated weekly to a mass accuracy of <0.1 ppm using a tuning solution from Agilent, which contains the following compounds: $C_2F_3O_2$, $C_6HF_9N_3O$, $C_{12}HF_{21}N_3O$, $C_{20}H_{18}F_{27}N_3O_8P_3$, and $C_{26}H_{18}F_{39}N_3O_8P_3$ with an *m/z* ranging between 112 and 1333. The instrument settings were optimized by tuning on a Suwannee River Fulvic Acid (SRFA) standard. Blanks (HPLC grade MeOH) were run at the beginning and the end of the day to monitor potential carry over from one sample to another and the instrument was flushed between samples using a mixture of water and methanol. The ion accumulation time (IAT) was varied to account for differences in C concentration between samples. One hundred and forty-four individual scans were averaged for each sample and internally calibrated using OM homologous series separated by 14 Da (-$CH_2$ groups). The mass measurement accuracy was <1 ppm for singly charged ions across a broad *m/z* range (i.e. $100 < m/z < 1100$). To further reduce cumulative errors, all sample peak lists for the entire dataset were aligned to each other prior to formula assignment to eliminate possible mass shifts that would impact formula assignment.

Putative chemical formulas were assigned using Formularity software[60]. Chemical formulas were assigned based on the following criteria: S/N > 7, and mass measurement error < 1 ppm, taking into consideration the presence of C, H, O, N, S and P and excluding other elements. Peaks with large mass ratios (*m/z* values > 500 Da) often have multiple possible candidate formulas. These peaks were assigned formulas through propagation of $CH_2$, O, and $H_2$ homologous series. Additionally, to ensure consistent choice of molecular formula when multiple formula candidates are found the following rules were implemented: the formula with the lowest error with the lowest number of heteroatoms was consistently picked and the assignment of one phosphorus atom required the presence of at least four oxygen atoms.

Rare peaks (occurring in only one of two replicate samples) and peaks <200 Da or >1,000 Da were removed prior to data analysis to increase confidence in the assignment of compounds to representative molecular classes. Compounds were then plotted on van Krevelen diagram based on their molar H:C ratios (*y*-axis) and molar O:C ratios (*x*-axis) [61]. Van Krevelen diagrams provide a means to visualize and compare the average properties of organic compounds and assign compounds to the major biochemical classes (e.g., lipid-, protein-, lignin-, carbohydrate-, and condensed aromatic-like). Biochemical compound classes were reported as relative abundance values based on counts of C, H, and O for the following H:C and O:C ranges: lipids ($0 < O:C \leq 0.3$ and $1.5 \leq H:C \leq 2.5$), unsaturated hydrocarbons ($0 \leq O:C \leq 0.125$ and $0.8 \leq H:C < 2.5$), proteins ($0.3 < O:C \leq 0.55$ and $1.5 \leq H:C \leq 2.3$), amino sugars ($0.55 < O:C \leq 0.7$ and $1.5 \leq H:C \leq 2.2$), lignin ($0.125 < O:C \leq 0.65$ and $0.8 \leq H:C < 1.5$), tannins ($0.65 < O:C \leq 1.1$ and $0.8 \leq H:C < 1.5$), and condensed hydrocarbons (aromatics; $0 \leq$ 200 $O:C \leq 0.95$ and $0.2 \leq H:C < 0.8$)[58]. Further, to disentangle essential elements and their relation to chemical class, each mass was then subsequently filtered with N>0, P>0 or S>0 and then plotted again on van Krevelen diagrams.

##### *1.4. Lipidomics Analysis*

Total lipid extracts (TLE) were analyzed as outlined in Kyle *et al.* (2017). Briefly, a Waters Acquity UPLC H class system interfaced with a Velos-ETD Orbitrap mass spectrometer was used for LC-MS/MS analyses. 10 µl of reconstituted sample was injected onto a Waters CSH column (3.0 mm x 150 mm x 1.7 µm particle size) and separated over a 34-minute gradient (mobile phase A: ACN/H2O (40:60) containing 10 mM ammonium acetate; mobile phase B: ACN/IPA (10:90) containing 10 mM ammonium acetate) at a flow rate of 250 µl minute$^{-1}$. TLEs were analyzed in both positive and negative modes, and lipids were fragmented using higher-energy collision dissociation and collision-induced dissociation.

#### 2. Supporting Tables and Figures

**SI Table 1.** Mineral Properties

| Property | Quartz | Kaolinite[a] | Ferrihydrite | Native Mineral[b] |
|---|---|---|---|---|
| Chemical Formula | $SiO_4$ | $Al_2Si_2O_5(OH)_4$ | $Fe(OH)_3$ | NA |
| Source of Mineral | Purchased from Sigma-Aldrich[c] | Purchased from Clay Minerals Society[c] | Synthesized in Lab[c] | Density Fractionated from Soil[c] |
| Initial C% | Negligible | Negligible | Negligible | 1.6 |
| BET Surface Area[d] ($m^2g^{-1}$) | 0.01-0.05[e] | 20.48 | 4.8 | 2.68 |
| Particle Size Range (µm) | 297-210 | Mostly < 2 | 297-210 | Not Determined |
| pH (1:1 m:v in 0.01 M $CaCl_2$) | 4.12 | 3.03 | 6.23 | Not Determined |
| Predicted Relative Charge Density | Very low | Low | High | Intermediate |
| Primary or Secondary Mineral? | Primary | Secondary | Secondary | NA |
| Hydroxyamine-Extractable Fe (mean ± SE, µg-1) | 8 ± 2 | 8.4 ± 0.4 | 4563 ± 1535 | Not Determined |
| Hydroxyamine-Extractable Al (mean ± SE, µg-1) | 1.5 ± 0.1 | 77 ± 14 | 587 ± 268 | Not Determined |

**a.** Kaolinite was used in a 50:50 mixture with quartz

**b.** Numerous parameters were not determined due to low quantities

**c.** See Supplemental Methods for detailed description of preparation

**d.** $N_2$ analysis gas

**e**. Quartz surface area was too low to measure using BET with $N_2$ gas. Estimated from the literature (Xu *et al.*, 2009, Mekonen *et al.*, 2013) and mesh size

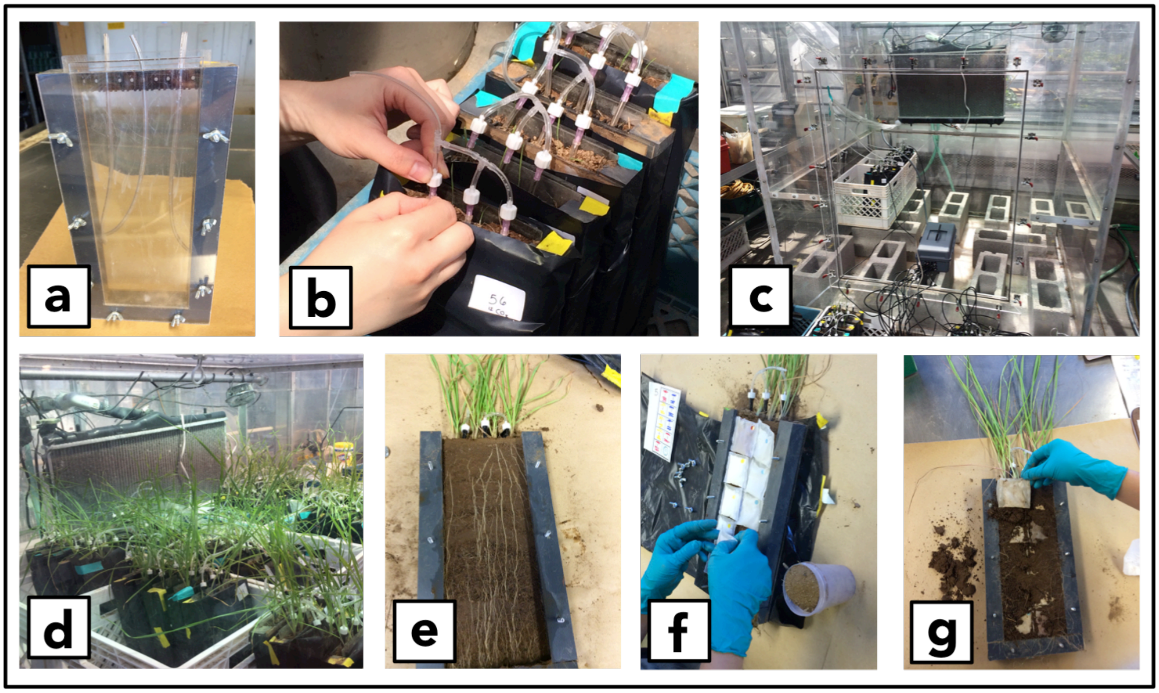


**SI Figure 1. Experimental design for mineral bags incubation in soil microcosms. Soil microcosms (a) were fitted with perforated tubing for automated watering and filled with soil (b). Soil microcosms were placed in an isotope labeling chamber (c) and *Avena barbata* seedlings were added to the rhizosphere treatment microcosms. Soil microcosms were tilted to 45° for 1 month (d), encouraging the *A. barbata* roots to grow along one face of the soil microcosm (e). Soil microcosms were opened and mineral bags were placed along the rhizosphere face for the rhizosphere treatment microcosms (f), or, for the bulk treatment with no plants, they were placed along the soil. Soil microcosms were placed back in the isotope labeling chamber (c), now sealed with added 99 atom% $^{13}CO_2$ maintained at 400 ppm with an atmospheric $^{13}CO_2$ concentration of ~ 10 atm% $^{13}C$. Soil microcosms were destructively harvested and mineral bags were recovered after 1, 2, and 2.5 months.**

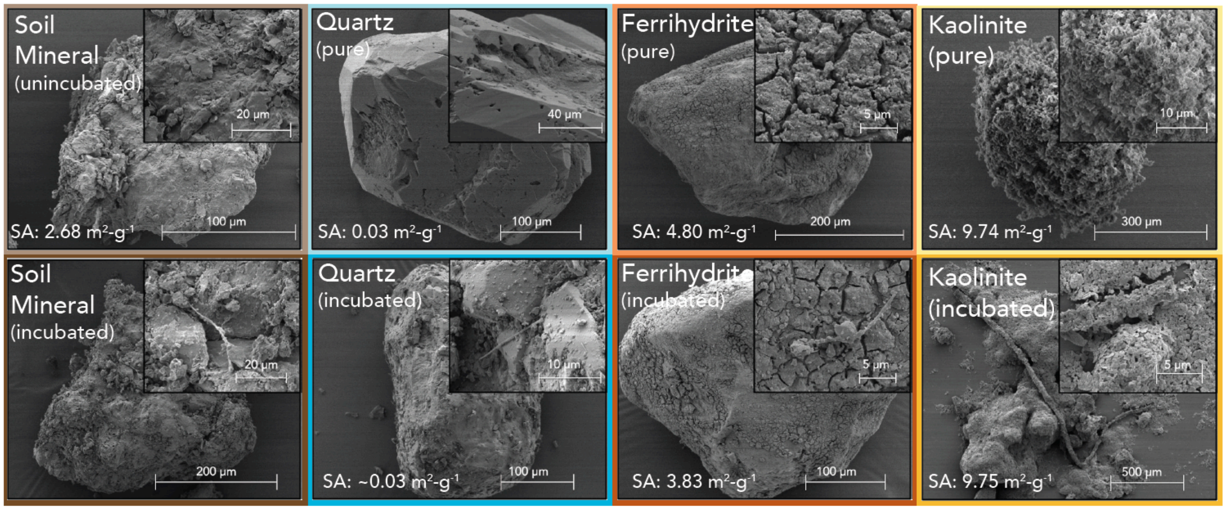


**SI Figure 2. SEM images (5 eV) of minerals before and after incubation in rhizosphere treatment soil microcosms. Pure minerals, shown in the top panels, are of minerals prior to incubation in soil microcosms. Minerals after incubation in the rhizosphere treatment soil microcosms are shown in the bottom panels. Each panel shows an overview of the mineral with an inset panel in the upper right hand corner showing a higher resolution image of the same mineral. Scale bars are in the bottom right of each panel and inset. SA stands for BET surface area, measured with $N_2$ gas.**

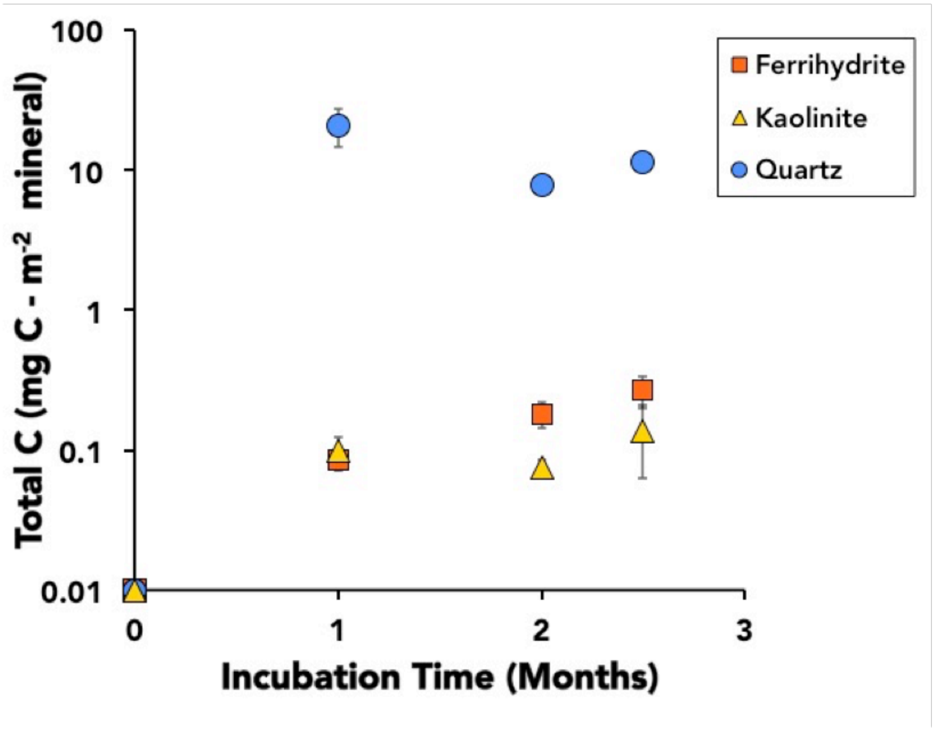


**SI Figure 3. Total C accumulation on pure minerals in rhizosphere treatment incubations, normalized to BET surface area. Quartz had the highest surface-area-normalized total C. All minerals began with no measurable C.**

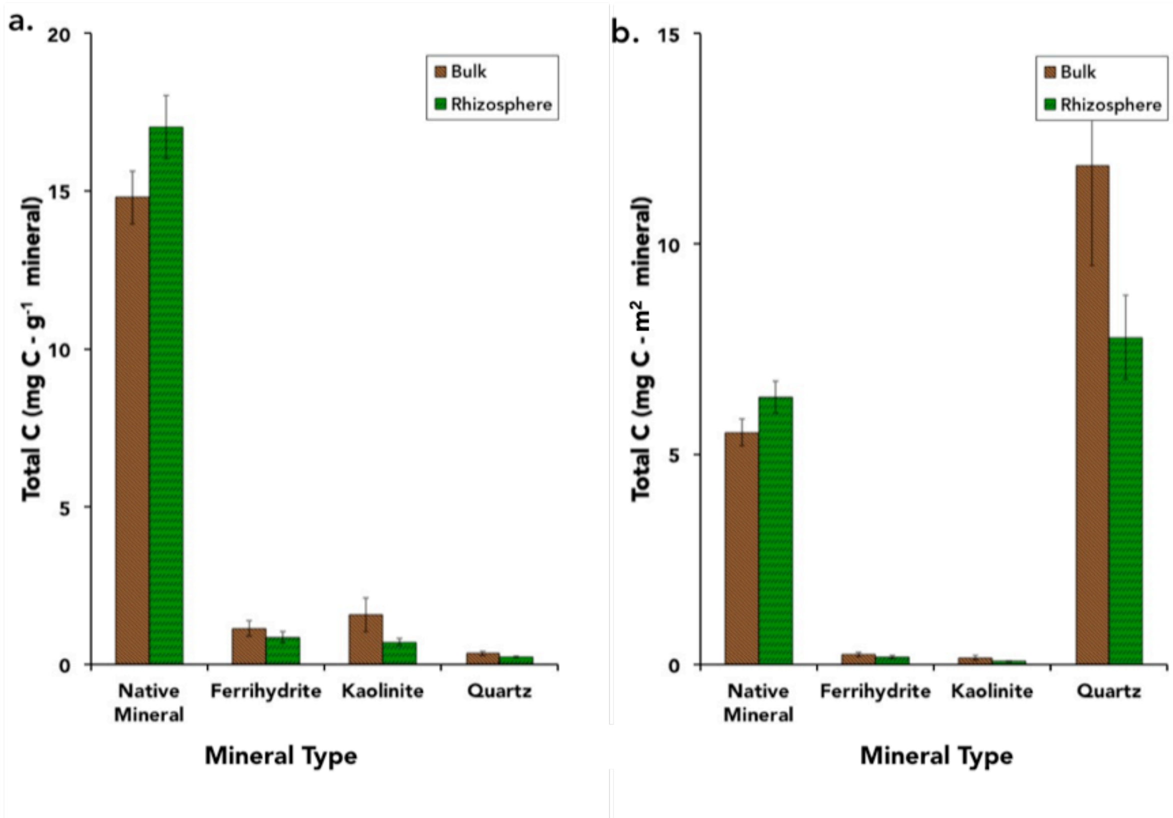


**SI Figure 4. Comparison of total C on minerals incubated in the rhizosphere and bulk treatments by mass (a) and normalized to surface area (b). The bulk soil treatment is shown in brown and the rhizosphere treatment in green.**

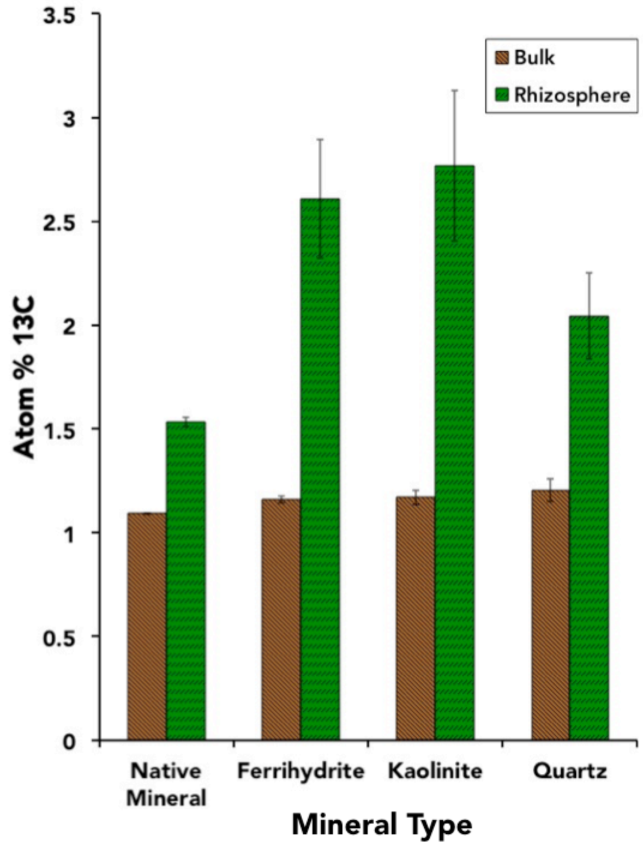


**SI Figure 5. Atom% $^{13}C$ of each mineral type at 2 months after incubation in soil microcosms.**

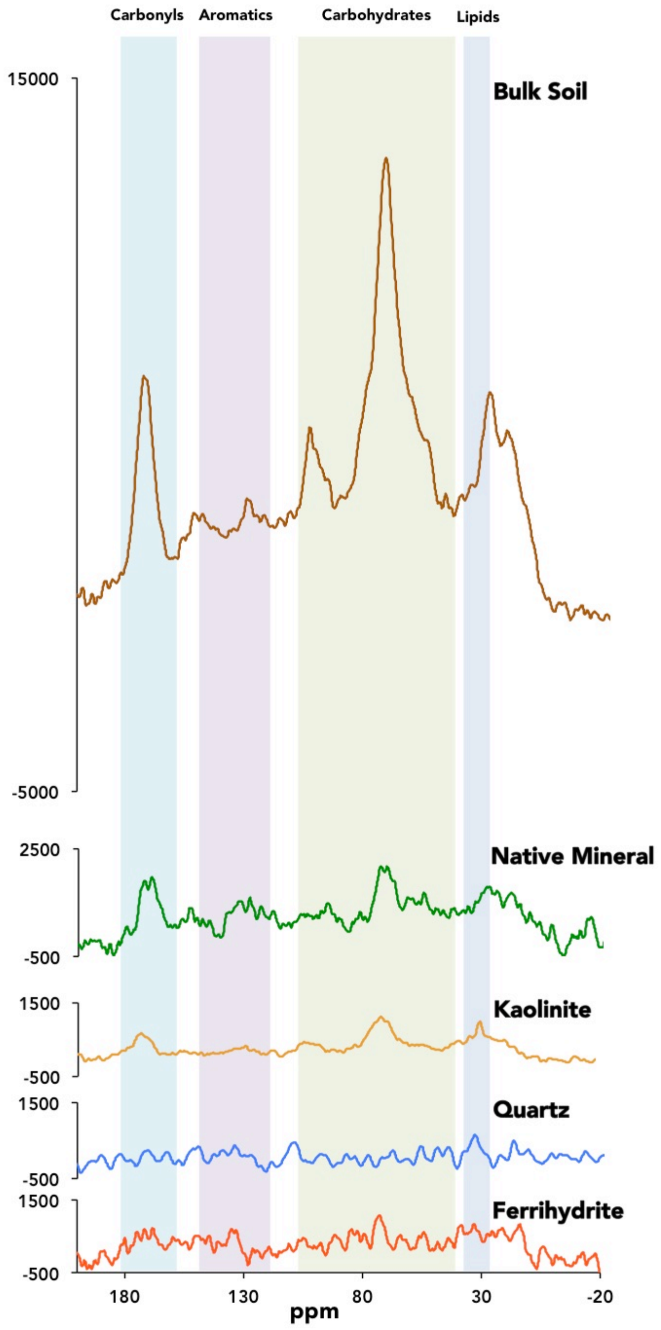


**SI Figure 6. $^{13}$C-NMR of all mineral types from the rhizosphere treatment compared with the bulk soil. The high iron content in the ferrihydrite caused interference, such that no clear spectra were obtained. Quartz, with a lower total C content by mass, also did not have well-resolved spectra.**

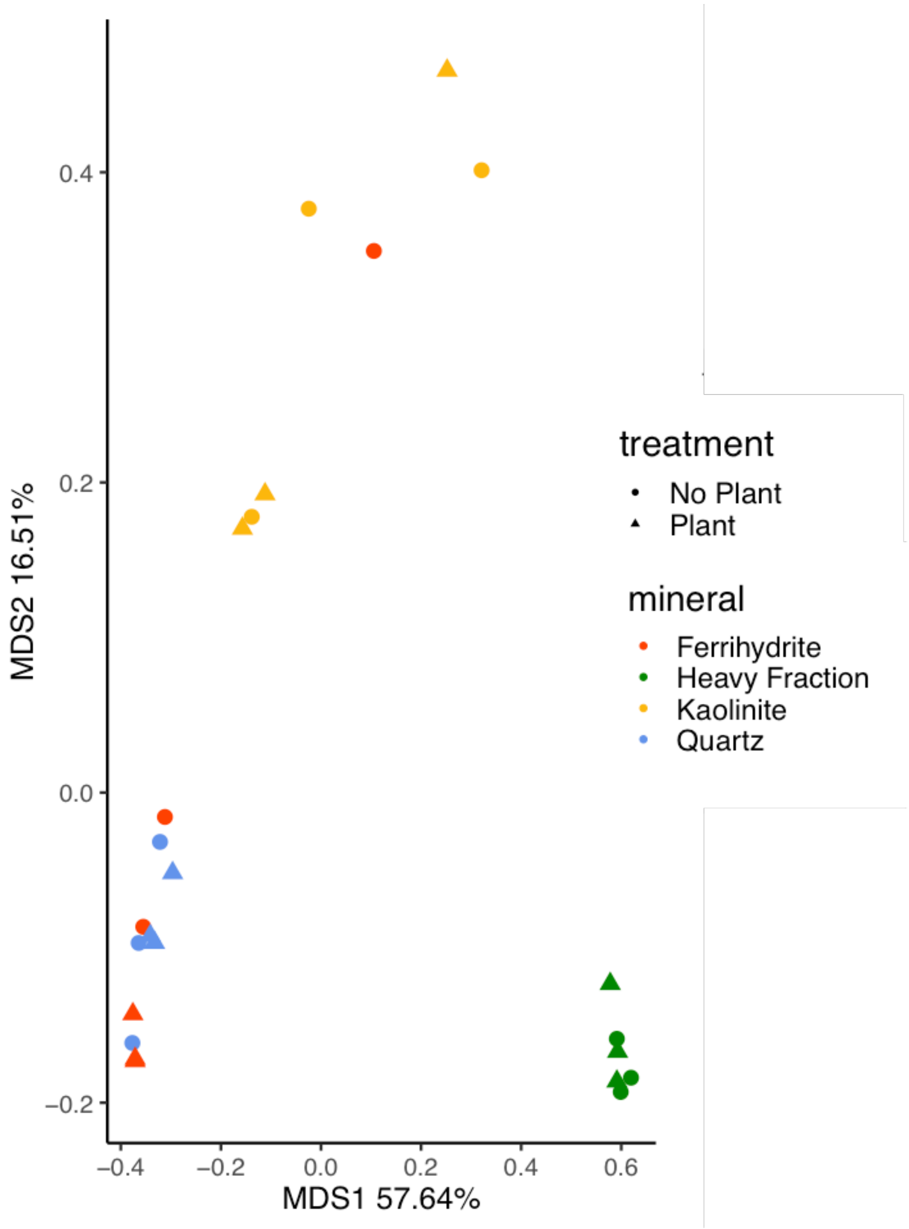


**SI Figure 7. PCoA of lipidomic data with Bray-Curtis shows clear clustering by mineral type, with the heavy fraction and the kaolinite in distinct clusters.**

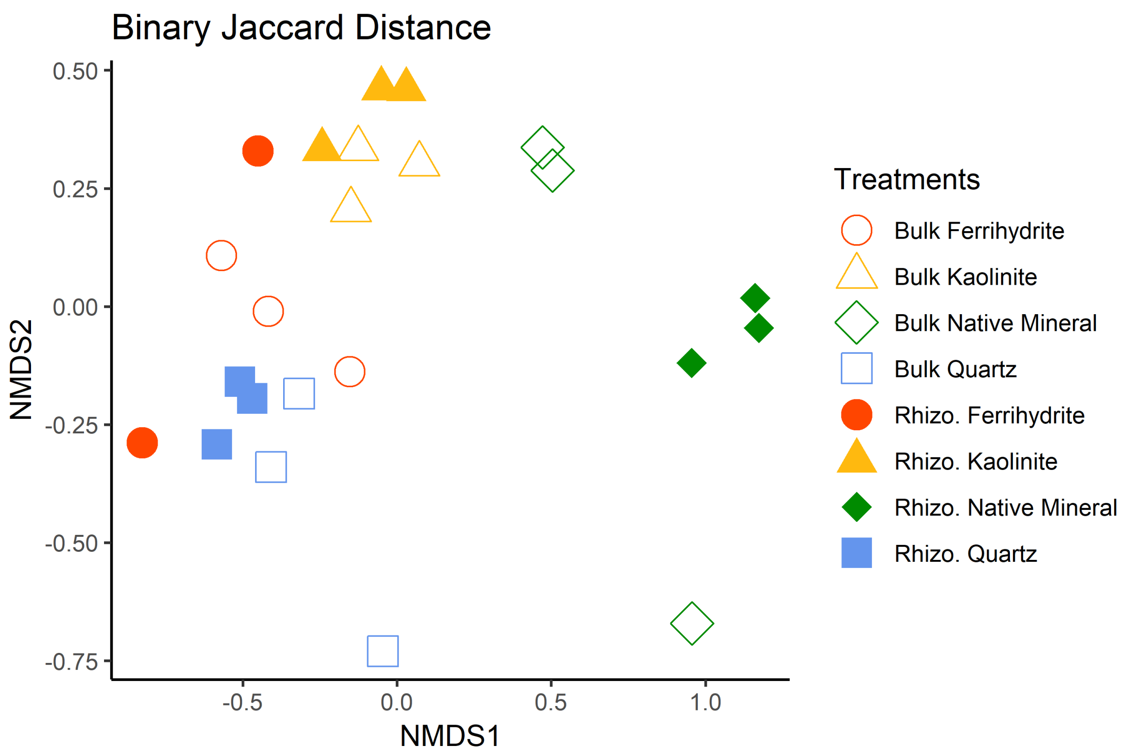


**SI Figure 8. NMDS of FTICR-MS data with Jaccard distance shows clustering by mineral type.**

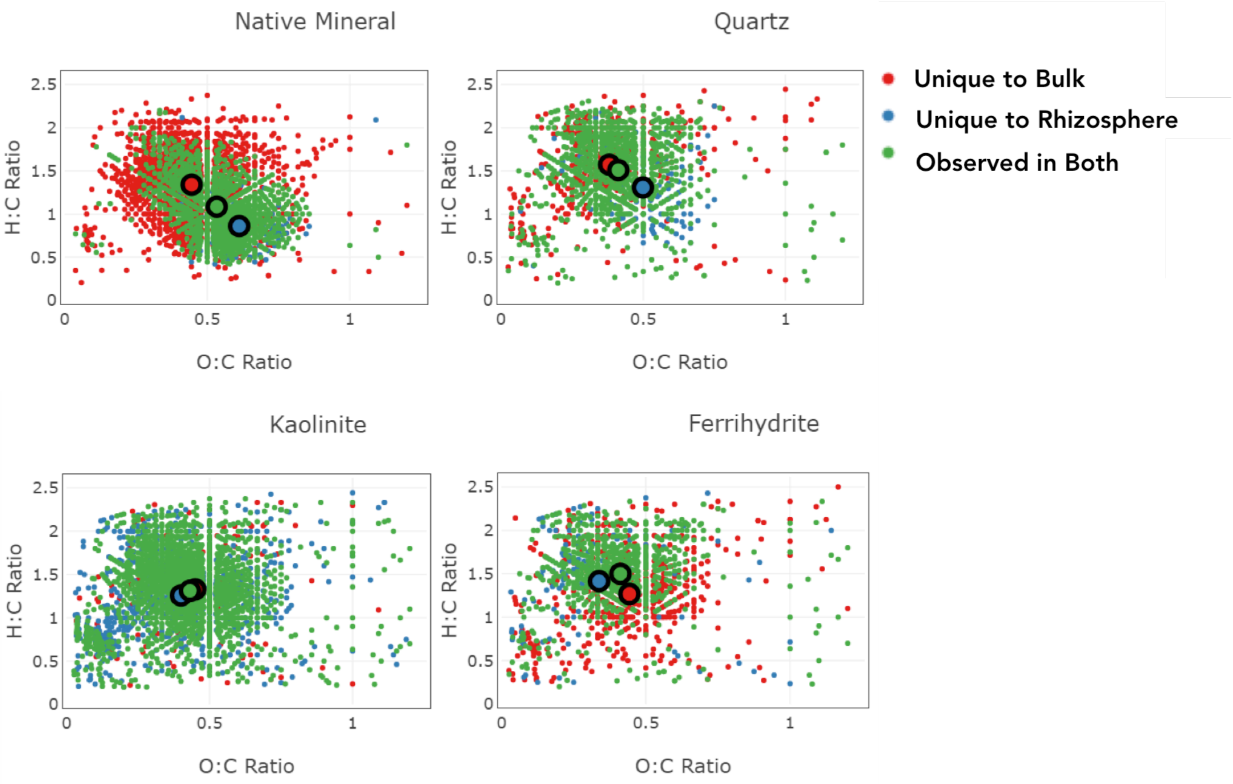


**SI Figure 9. Van Krevelen diagrams of FTICR-MS data, with a graph for each mineral type. Compounds unique to the bulk are in red, unique to the rhizosphere in blue, and observed in both are green. The center of mass is shown for each.**

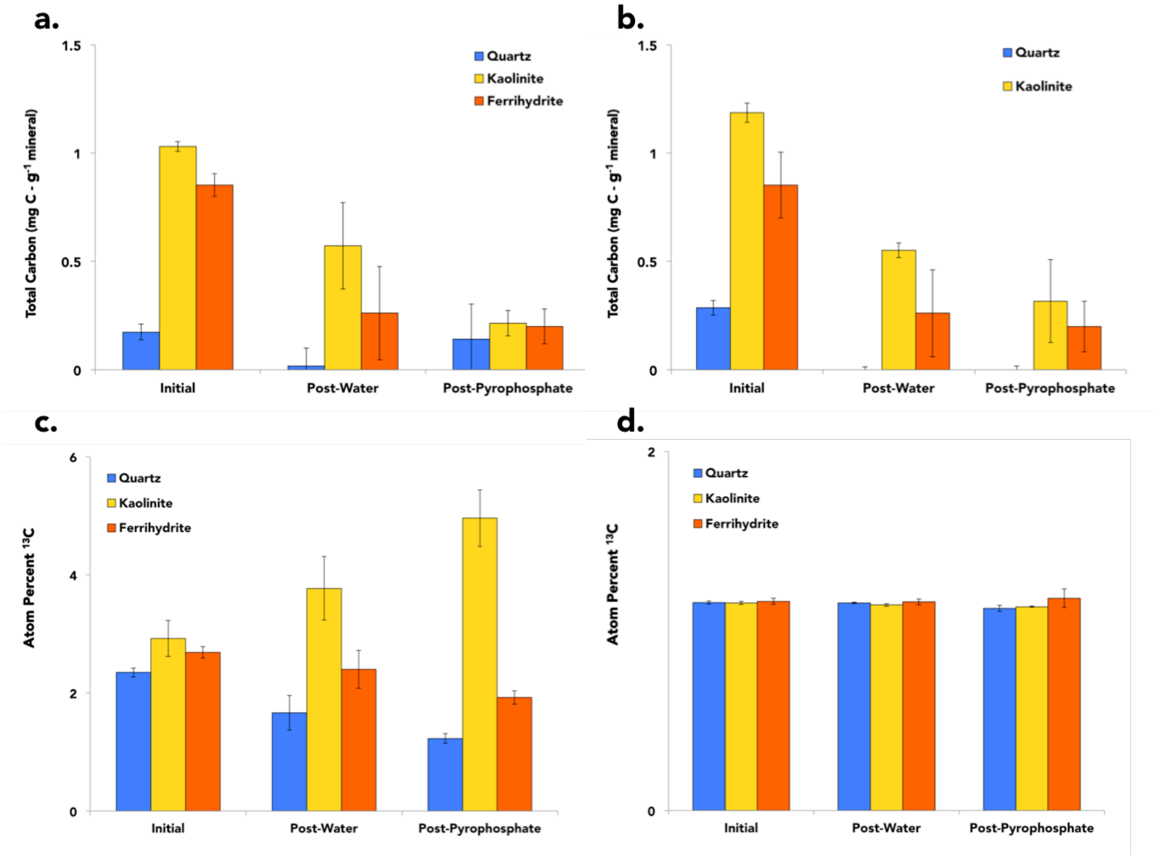


**SI Figure 10. Total C after sequential extractions of mineral-associated soil organic matter (SOM) for the rhizosphere treatment (a) and bulk soil (b), compared to initial total C. Changes in atom% $^{13}C$ after sequential extractions for the rhizosphere treatment (c) and bulk soil (d), compared to initial atom% $^{13}C$.**
